# Spatial pattern analysis of line-segment data in ecology

**DOI:** 10.1101/2021.01.18.427207

**Authors:** Luke A. Yates, Barry W. Brook, Jessie C. Buettel

## Abstract

1. The spatial analysis of linear features (lines and curves) is a challenging and rarely attempted problem in ecology. Existing methods are typically expressed in abstract mathematical formalism, making it difficult to assess their relevance and transferability into an ecological setting. A set of concrete and accessible tools is needed.
2. We develop a new method to analyse the spatial patterning of line-segment data. It is based on a generalisation of Ripley’s *K*-function and includes an analogue of the transformed *L*-function, together with estimators and theoretical expectation values. We introduce a class of line-segment processes, related to the Boolean model, which we use in conjunction with Monte-Carlo methods and information criteria to generate and compare candidate models. We demonstrate the utility of our method using fallen tree (dead log) data collected from two one-hectare Australian tall eucalypt forest plots.
3. Comparing six line-segment models, we find for both plots that the distribution of fallen logs is best explained by plot-level spatial heterogeneity. The use of non-uniform distributions to model dead-log orientation on the forest floor improves model performance in one of the two sites. Our case study highlights the challenges of model comparison in spatial-pattern analysis, where Monte-Carlo approaches based on the discrepancy of simulated summary functions can generate a different ranking of models than that of information criteria.
4. These methods are of a general nature and are applicable to any line-segment data. In the context of forest ecology, the integration of fallen logs as linear structural features in a landscape with the point locations of living trees, and a quantification of their interactions, will yield new insights into the functional and structural role of tree fall in forest communities and their enduring post-mortem ecological legacy as spatially distributed decomposing logs.

## Introduction

Forest ecologists often use ‘snap-shot’ patterns of individual trees, taken from measurements of their spatial positions, to recover information of past processes, building an ‘ecological archive’ of the forest community (Wiegand et al., 2003). Detailed information obtained using this approach is based on the location and attributes of the living trees, and typically analysed using spatial point-pattern analysis (SPPA e.g., (Law et al., 2009; Perry et al., 2006)). Spatial-pattern analysis complements spatially explicit individual-based models for uncovering the relationship between pattern and process (Pommerening et al., 2011). The stochastic processes captured therein can be arbitrarily complex, modelling a range of ecological interactions. For this purpose, there exists a vast array of well-studied model classes including cluster, hard-core, Gibbs (e.g., (Illian, 2008, ch.6)). While the list of applications is extensive, they have been largely restricted to problems that can be represented as a point pattern; that is, as a set of dimensionless points within a spatial domain.

However, in forest ecology, there is increasing awareness and recognition of the critical role that fallen/dead wood play in the structure and function of forest ecosystems (e.g., decomposition, structural complexity, mortality, recruitment, e.g., (Buettel et al., 2017)). However, despite this, the dead elements (e.g., decaying logs, fallen trees) are rarely measured or analysed as linear features in an SPPA framework (Buettel et al., 2018). There has been some success in modifying spatial point-pattern statistics, like Ripley’s *K*, to deal with points distributed along linear networks (Spooner et al., 2004; Baddeley et al., 2020). There are also examples of modelling lines and points separately and then testing these for interactions, for instance in geological studies of ore seams (e.g., Mardia (1972)), see also (Stoyan and Ohser, 1982). Or, more commonly, only the point location of the base of the fallen log is measured and analysed using visual techniques like ‘rose plots’ or using point-based SPPA methods (Oberle et al., 2015; Rouvinen and Kuuluvainen, 2001). But unlike the widespread utility of SPPA of zero-dimensional (point) locations, there remains little statistical development for one-dimensional (linear) features within the same framework, aside from their mapping and visualisation (Buettel et al., 2018). Such utility would not only improve our knowledge of a forest’s past and comprehension of its present, but also our capacity to predict — and possibly manage — its future.

A typical workflow in ecological SPPA includes the specification of a null hypothesis, the selection of appropriate summary statistics, and computation of the associated *p*-values, together with accompanying plots such as (Monte-Carlo) simulation envelopes (Law et al., 2009; Perry et al., 2006). Common summary statistics include first-order — expected density — statistics such as distance to nearest neighbour, and second-order — accounting for spatial (co)variance — statistics such as Ripley’s *K* and its transformed variants (Wiegand et al., 2013). Statistical methods for point-pattern analysis are at a mature stage of development (Velázquez et al., 2016), and many of the most popular techniques have been robustly implemented in user-friendly software such as the R package spatstat (R Core Team, 2020; Baddeley et al., 2015). When implemented, the SPPA process can be iterated, often starting with the null model of complete spatial randomness (CSR) before advancing to more sophisticated hypotheses which can include tests of spatial heterogeneity, dependence on attributes (marks) of each point, or complex point-wise (paired) interactions.

In contrast, existing methods for the analysis of spatially distributed linear features (fibres) remain couched in the mathematical formalism of stochastic geometry (Chiu et al., 2013, Chap.8). The abstract presentation of these methods makes it difficult to translate them into ecological contexts (e.g., see the second-order summary functions presented in (Stoyan, 1983; Chiu et al., 2013, p.319). Furthermore, the underlying definition of these statistics is based on the expected distribution of the line segments (or, ‘fibres’) within the neighborhood of a random point on the fibre system; this is less suitable for ecological questions where the distribution within the neighborhood of an entire (random) fibre is more meaningful. Some initial progress towards a summary function for fibre processes in an ecological context is given by Fortin and Dale (2005, p.62).

In this paper, we used pre-measured and mapped fallen-log data from two one-hectare plots in southern Australian tall eucalypt forests as a case study to showcase the development of a new statistical method, based on a natural extension of the second-order summary function, Ripley’s *K*. We model and represent the treefall data as a two-dimensional (finite-length) line-segment pattern, and introduce a class of stochastic line-segment processes, demonstrating how to apply both Monte-Carlo techniques and information-theoretic model scores to compare candidate models. Expressed in concrete and accessible notation, we define both estimators and theoretical expectation values for the new summary function; these include edge-correction terms (measurement plots are bounded), and an analogue of the pair correlation function (PCF). These methods are applicable to any line-segment data (i.e., not restricted to treefall) and they provide a foundation for the development of extended techniques such as bivariate and inhomogeneous generalisations, as well as the unification of line-segment and point data into a single analytic framework.

## Method Development

### Background: Ripley’s *K*

Two-dimensional point-pattern analysis is often summarised using the second-order summary function, Ripley’s K (*K*(*r*), Ripley (1977)). Assuming a complete census of points within a fixed, two-dimensional observation window, *W*, the starting point is the selection of a second-order summary statistic, *S*(*r*), which is taken to be the expected number of points within a circle of radius *r*, centred on a randomly chosen point. The associated *K*-function is defined by^1^

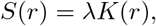

where *λ* is the density of points per unit area. An unbiased estimator for *K*(*r*) is given by

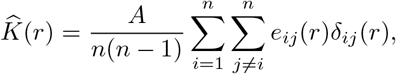

where *n* is the number of points, *A* is the area of the observation window and *e_ij_*(*r*) is the edge correction term (discussed below). The final term, *δ_ij_* (*r*), is an indicator function taking the value 1 if *d_ij_* ≤ *r* and 0 otherwise, where *d_ij_* is the distance between the points *i* and *j*.

The term preceding the summation standardises the definition of 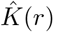, making it possible to compare point patterns with different numbers of points and different observation windows (Baddeley et al., 2015); the summation term itself may be viewed as a bias-corrected estimate of *S*(*r*).

### A Summary Function for Line-Segment Patterns

To develop a summary function analogous to Ripley’s *K* for fallen-log (herein, line-segment) data, we must decide which of the available attributes should be used. Candidate line-segment attributes include: length, number, number of intersections (where the logs ‘cross over’) as well as quantities derived from the directional data such as average angle, angular distribution, and vector operations such as inner products. The sample area can be defined as the region within a fixed distance *r* from either, a) any point along the length of a random segment, or b) from a single random point on a random segment. The latter defines a circular region, most commonly applied and referenced in scientific fields such as statistics and stereology (Chiu et al., 2013). In these fields, the selected attribute is the net length of segments (within a given neighbourhood) since the total number of lines and the number of intersections may be identified as associated point patterns. Determining the relationship between various classes of stochastic geometric patterns in two-(*resp.* three-) dimensions and their associated (cross-sectional) patterns in one-(*resp.* two-) dimensions are key results in stereology (Chiu et al., 2013).

In the context of this ecologically focused paper, where each line segment is itself the natural unit of observation (a fallen log), we take a complete segment as the representative sample ‘point’ and base the corresponding statistic on what is observed within its entire neighbourhood (as suggested by Fortin and Dale (2005)). Therefore, we define *S*(*r*) to be the expected net length of line segments found within a distance *r* of a randomly chosen segment, introducing the following estimator,

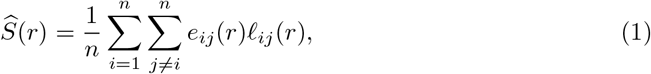

where *n* is the total number of line segments, *e_ij_*(*r*) is the edge correction term and *ℓ_ij_*(*r*) is the length of segment *j* lying within distance *r* of segment *i*. Before defining the associated *K*-function, let us first consider the theoretical expectation value, *S_theo_*(*r*), of a line-segment pattern that is generated by a process of complete spatial randomness (CSR). For line-segment processes, there is not a unique definition of CSR, since the expected value depends upon the distribution of segment lengths (Hanisch et al., 1985). Here we sum over the empirical set of segment lengths, to define theoretical expectation value for CSR as,

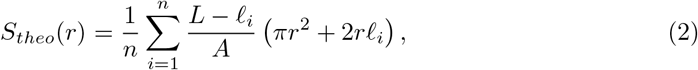

where *ℓ_i_* is the length of segment *i* and *L* = *∑_i_ ℓ_i_* is total length of all segments. The term *πr*^2^ + 2*rℓ_i_* is the sample area surrounding segment *i* and 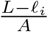 is the (homogeneous) density of line segments, calculated by excluding the length of the sample segment.

### Edge Corrections

Appropriately accounting for edge effects is a standard requirement in spatial statistics. For summary statistics such as the *K*-function, the edge-correction term adjusts for the bias that arises from estimation within a finite observation window (Illian, 2008, chap 4). In point-pattern analysis, given a circle of distance *d_ij_* ≤ *r*, centered on point *i*, the edge correction *e_ij_*(*r*) is generally taken to be the reciprocal of the proportion of that circle’s circumference within *W*. The choice of method for calculating this term amounts to a trade-off between the accuracy and the computational demands of the implemented algorithm. For the case at hand, where the sample area is non-circular and an (unbiased) edge correction term would vary continuously with *r*, we are unable to exploit the geometric shortcuts that have been developed for point patterns (Goreaud and Pélissier, 2006). We use instead a simplified method, as suggested by Dale2005, for which the correction term, *e_ij_*(*r*) = *e_i_*(*r*), depends only on the reference segment *i* and the distance variable *r*. Given a sample area whose perimeter is distance *r* from the closest point on line segment *i*, we define the corresponding edge correction to be the reciprocal of proportion of that sample’s *area* within *W* (Figure 1A).

**Figure 1:**
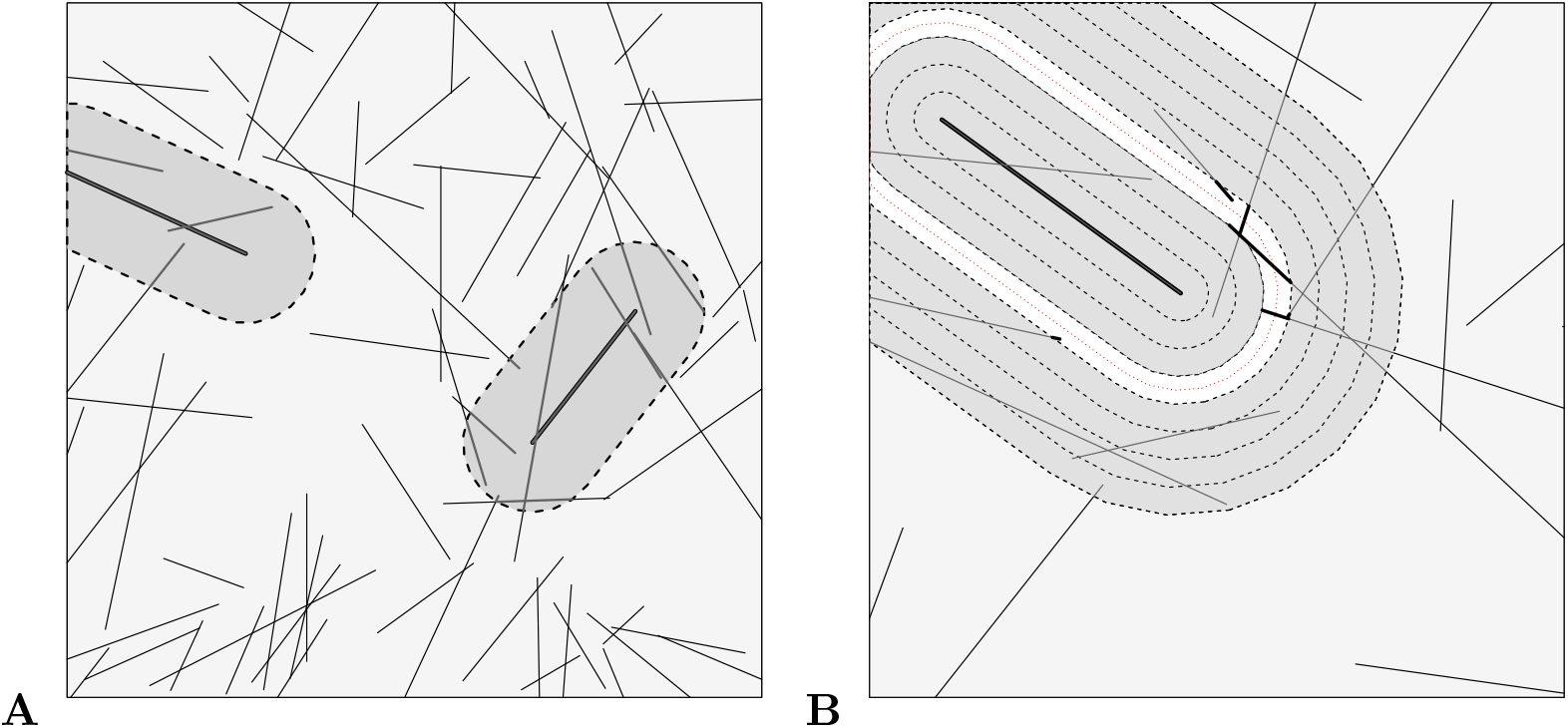
Edge correction examples. A) The sample area on the left is clipped by the boundary; the corresponding edge correction is 1.22. The other area is completely within the site boundary; its edge correction is 1. B) Illustration of the concentric ‘rings’ method for improved edge correction as well as a numerical approach to estimate the pair correlation function.

If increased precision of the estimated edge correction term is required, a concentric ‘rings’ approach can be used, in which sub-segments along each segment are binned into discrete intervals surrounding the reference segment *i* (Figure 1B). The contribution of each sub-segment to the summary function is weighted according to the proportion of the perimeter of the corresponding bin within *W*. This approach is more computationally intensive than the proposed method described above, and can be viewed as a discrete approximation to the continuous approach of evaluating line integrals along each segment.

### Transformations of *S(r)*

The specific dependence of *S_theo_*(*r*) on the (empirical) distribution of log lengths makes it difficult to define a standardised form of *K*(*r*). As a first step, we rearrange the summation in (2), so as to express *S_theo_*(*r*) in terms of the mean and variance of the segment lengths,

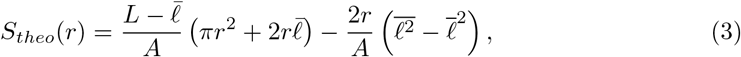

where 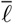 denotes the mean of *ℓ*, etc. It would be convenient to transform *S_theo_*(*r*) into the familiar form *K_theo_*(*r*) = *πr*^2^ which could then be linearised using the variance-stabilising transform 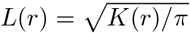 (Besag, 1977). The first part is easily achieved via addition and multiplication of the appropriate terms, leading to

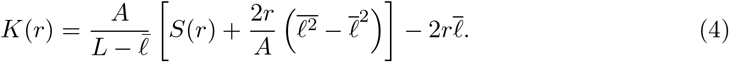

However, in applying this transformation to the estimator 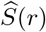 given in equation (1), the resultant function 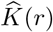 is no longer positive definite, which in turn prohibits linearisation via the square-root transform. This can be addressed using a simple complete-the-square transformation such that 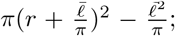 is expressed as 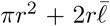 this suggests, as an intermediate step, the definition,

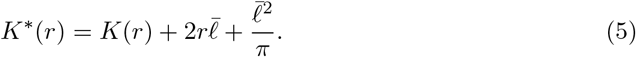

It follows that *K*^*^(*r*) is a positive-valued, monotonic increasing transformation of *S*(*r*), permitting the linearisation,

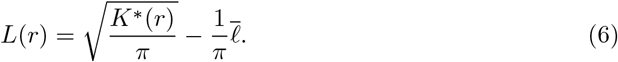

The corresponding theoretical curve assumes the familiar form from point-pattern analysis,

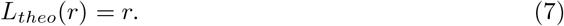

In practice, it is usually *L*(*r*), rather than *K*(*r*), that is employed as a summary function for point patterns as it is more easily visualised and has improved statistical properties Diggle 2013; this same reasoning suggests its preferential use for line-segment analysis.

### Pair correlation function

The pair-correlation function (PCF), *g*(*r*), can be defined in terms of the *K*-function,

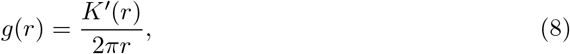

where *K*^′^(*r*) is the derivative of *K* with respect to *r*. Equivalently, *g*(*r*) can be defined in terms of the probability of finding two ‘points’ each within an ‘infinitesimal’ region of space separated by distance *r*. In either case, the function must be estimated by either binning, kernel smoothing, or differentiating splines that have been fitted to 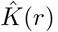 (Baddeley et al., 2015).

The relative merits of the PCF with respect to the *K*-function have be extensively debated in the literature (Illian, 2008). It is commonly stated that *K*(*r*) is the most appropriate for statistical inference, provided its variance has been stabilised via transformation to *L*(*r*) (Law et al., 2009; Baddeley et al., 2015); however, this view has been challenged by simulation studies (Wiegand et al., 2013). The PCF, which can be viewed as a probability density, gives critical information about the scale of spatial structure which might otherwise be obscured by the cumulative *K*-function (Wiegand and A. Moloney, 2004; Velázquez et al., 2016). However, due to its cumulative nature, *K*(*r*) performs favourably for small data sets where *g*(*r*) is known to be noisy (Diggle, 2013, p.71). These comments apply equally in the context of fibre processes, although the general problem of estimating the summary functions is substantially more difficult (Stoyan, 1985; Chiu et al., 2013).

Here we use (8) to evaluate the PCF by differentiation of smoothing splines fitted to our estimate of *K*(*r*). We found that *g*(*r*) was highly variable for small values of *r*, even after the imposition of additional constraints at the origin following Baddeley et al. (2015). Given this, we note that the construction of concentric ‘rings’, discussed above in the context of edge corrections, could be employed to generate kernel estimates of *g*(*r*), in which adaptive bandwidths can be used to address variability issues (Illian, 2008, p.232).

## Line Segment Models and Model Selection

### The Boolean Model

In the simplest case, a line-segment model can be viewed as a natural extension of a Poisson point-process model. Formally, these models belong to the class of *Boolean models* which have been studied extensively within the mathematical literature (Hanisch et al., 1985; Chiu et al., 2013, Chap.8). We present one instance of these models, which we call a *line segment model*. For a given two-dimensional observation window *W*, we express this model in terms of the following three functions:

- *λ* = *λ*(*x, y*), the intensity, possibly inhomogeneous, of a Poisson point process which generates one end-point for each segment;
- *R*(*θ*), a circular distribution function of segment angles with respect to a fixed axis; and
- *F* (*ℓ*), a distribution function of segment lengths.

It is important to differentiate between directed, 0 ≥ *θ* ≥ 2*π*, and undirected, 0 ≥ *θ* ≥ *π*, line-segment processes. We use the former in this case study, since the base of a tree provides a natural orientation. The set of all bases are the end-points for the Poisson point process described by *λ*.

### Model Comparison

We present two different approaches to compare candidate models. In the first approach, Monte-Carlo simulations are used to compute a discrepancy with respect to the summary function *L*(*r*). This permits model comparison in the absence of an analytic likelihood, such as when using kernel-density estimation or minimum-contrast estimation. The second approach assumes that all models are parametric, using the maximum likelihood to compute the Akaike Information Criterion (AIC).

#### Monte-Carlo discrepancy

For each model, the *L*-function (6) is computed for both the observed data and a set of model simulations. The goodness-of-fit is measured by the discrepancy *D* = *D*(*R*), defined as the square of the area between the empirical summary function 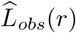 and the average of the simulated summary functions 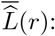:

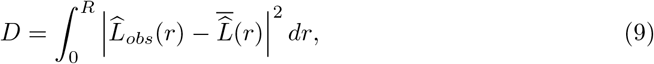

where *R* is the maximum neighbourhood distance under consideration. Smaller values of *D* indicate a better fit. *D* is equivalent to the Diggle-Cressie-Loosmore-Ford (DCLF) test statistic, used to define a Monte-Carlo test based on the rank of the empirical *D*-value amongst those of model simulations (Illian, 2008, p.455). (Note, 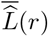 *includes* the empirical *L*-function, to preserve symmetry, when used to compute a Monte-Carlo *p*-value, but excludes it when used for a goodness-of-fit measure.) *D* is also closely related to the Cramér-von Mises test statistic, and the least-squares discrepancy used in minimum contrast estimation (Baddeley et al., 2015, p.381;484).

#### Information criterion

AIC is an estimate of expected, relative Kullback-Leibler discrepancy, a measure of out-of-sample model performance related to the information lost when an approximating (simpler) model is used in place of the true, but typically unknown, data-generating model (Akaike, 1973). It is computed by adding a bias correction to the within-sample (maximum) log-likelihood of the data. Assuming a large sample size, the criterion is given by:

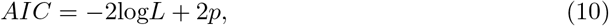

where *L* is the likelihood of the data and *p* is the number of estimated parameters; smaller AIC values indicate better model performance. For Poisson models of point-pattern intensity, parameters may be estimated using a Berman-Turner device (Berman and Turner, 1992). This allows the point process to be approximated by a generalised linear model, where a numerical quadrature scheme encodes the spatial observations as discrete Poisson-distributed events. The log-likelihood of a line segment model can be expressed as

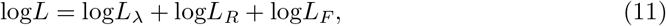

where *L_λ_* is the likelihood of observing the segment end-points, computed from the residual deviance of the generalised linear model (Berman and Turner, 1992, eq (3.2)), *L_R_* is the likelihood of the angular data, and *L_F_* is the likelihood of the length data.

#### Comparison of methods

Information-theoretic approaches provide a rigorous foundation for model comparison. When models are parametric and estimated using maximum likelihood, AIC is a convenient and rapidly computed model score, which avoids the computational cost of evaluating the summary function for each simulation. On the other hand, the summary functions provide a different perspective on the model fit. By construction, the *L*-function summarises information about the density of segments in neighbourhoods of different sizes. The discrepancy *D* can be computed for a chosen value of *R*, allowing models to be compared at a specific distance of interest; AIC does not allow distance-specific comparisons between models.

The problem with using a goodness-of-fit measure such as *D* to compare and ultimately select amongst candidate models is that this will often lead to over-fitting (i.e., low-bias but high-variance models). Information criteria are usually preferred for model selection because they estimate out-of-sample performance, thus achieving as trade-off between goodness-of-fit and generalisation to new data (Burnham and Anderson, 2002). In practice, due to analytical constraints, different candidates within the model set may be fit using different estimation methods, making model comparison a difficult and nuanced task, for which both *D* and AIC can provide useful insight.

Monte-Carlo methods in point-pattern analysis are often associated with null-hypothesis testing. Envelope tests, carefully constructed to avoid the problems of multiple comparisons on the interpretation of *p*-values, allow excursion of the empirical summary function beyond the envelope to be interpreted a statistically significant (Baddeley et al., 2014). Two-stage Monte-Carlo methods allow valid testing when the null hypothesis is composite (Baddeley et al., 2017). Existing hypothesis tests for point patterns, based on the summary functions of simulated realisations of a null model, are easily applied to line segment models, although computational cost could make two-stage tests prohibitive.

## Case Study

### Data Sets

In 2016, 12 of the 48 plots from the Ausplots Forest Monitoring network (see Wood et al. (2015) for further details on the locations, establishment and characteristics of the 48 plots) were re-visited and additional measurements of the coarse-woody debris were taken. For all ‘large’ fallen logs (≥ 5*m* length and ≥ 20*cm* DBH) the spatial location (precise coordinates, on a two-dimensional grid) of the base of the log, length, diameter-at-breast-height (DBH), and angular direction were recorded. The DBH of the logs was obtained by measuring the half circumference of the stem at 1.3 m from the base of where the uprooted or broken tree would have emerged above ground (or at the top, if the log was shorter than breast height) with a DBH tape. We used two of the 12 measured plots as case studies to demonstrate the utility of our method. One of the plots was located in Tasmania (T-SX, latitude/longitude location: −42.8118, 146.6083) and the other, in Western Australia (W-FR, location: −34.8247, 116.8734) (Figure 3). The dominant tree genus within these plots is *Eucalyptus*, with a rainforest and mixed-shrub understorey, respectively. Slope data, used to create digital-terrain surfaces for each plot, were collected in the field at a resolution of 10 m for all 12 plots, as per the method described in (Buettel et al., 2018) (see supplementary materials for slope and elevation plots).

### Candidate models

We selected six representative line-segment models, differentiated by two types of intensity functions, *λ*, and three types of angular-distribution functions, *R*(*θ*). For all models we fit a log-normal distribution, *F*(*ℓ*), to the set of fallen-log (segment) lengths within each plot, omitting those which intercept the boundary as their full lengths were not recorded. An overview of the model functions is given in Table 1.

**Table 1.** Description of model functions.

|  |  |  |
| --- | --- | --- |
| $\lambda$ | *Homogeneous* | Constant intensity, $\hat{\lambda} = A/n$ |
| | *Inhomogeneous*<sup>†</sup> | Log-quadratic function of the Cartesian coordinates,<br>$\log \hat{\lambda} = \beta_0 + \beta_1 x + \beta_2 y + \beta_3 x^2 + \beta_4 y^2 + \beta_5 xy$ |
| $R(\theta)$ | *Uniform* | Uniform angular distribution |
|  | *Re-sampled* | Angles re-sampled with replacement |
|  | *Local*<sup>†</sup> | Circular distribution fit to the set of relative angles between the direction of each log the local gradient at the log base<sup>§</sup> |
| $F(\ell)$ | *Log-normal*<sup>†</sup> | Log-normal distribution fit to the set of logs that do not intersect the boundary |

<sup>†</sup>Parameters estimated using maximum likelihood
<sup>§</sup> See supplementary materials for details.

The candidate models can be used to investigate the dependence of the pattern of fallen logs on inter-tree spacing, tree heights (fallen-log lengths) and the probability of a tree falling in a given direction. The log-quadratic intensity models account for spatial heterogeneity of the location of fallen-log bases, the re-sampling of fallen-log orientations captures existing processes that determine direction of treefall, such as prevailing wind (Ulanova, 2000), and the local gradient-fitted distribution of angles is based on the known result that trees are more likely to fall downhill (Buettel et al., 2018). Finally, the homogeneous intensity paired with the uniform distribution of orientations acts as a null model of complete spatial randomness.

### Analysis

For each model, we compute the discrepancy *D* = *D*(*R*) (9), setting *R* = 25*m* and using 99 Monte-Carlo simulations, and the AIC score (10) (not available for re-sampled models). The resulting values of these statistics for each of the model fits are presented in the subtitles of the corresponding plots in Fig 2.

**Figure 2:**
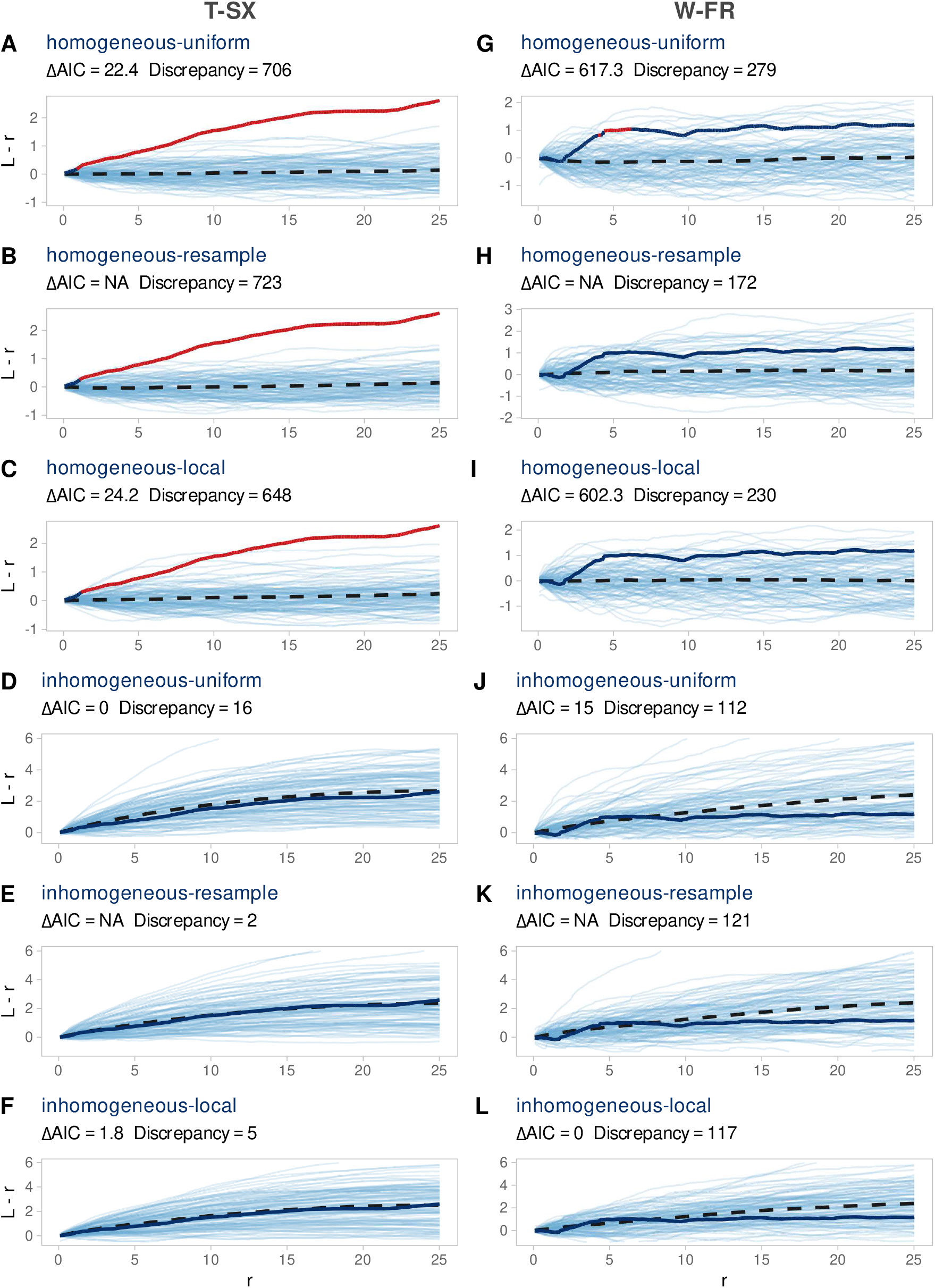
Centred summary functions *L*(*r*) − *r* for both sites. The light blue lines depict the 99 Monte Carlo simulations, the solid line is the empirical curve, and the dashed line is the mean value of all *L*(*r*) − *r*. Viewed as a null hypothesis test for each model, the progress of the Monte-Carlo *p*-value associated with the rank of DCLF test statistic (9), where *R* = *r*, is indicated by the colour of the empirical curve: red if *p* ≤ 0.05, otherwise dark blue. T-SX (Tasmania-Styx) and W-FR (Western Australia-Frankland) are the names of the two eucalypt-forest plots analysed herein.

**Figure 3:**
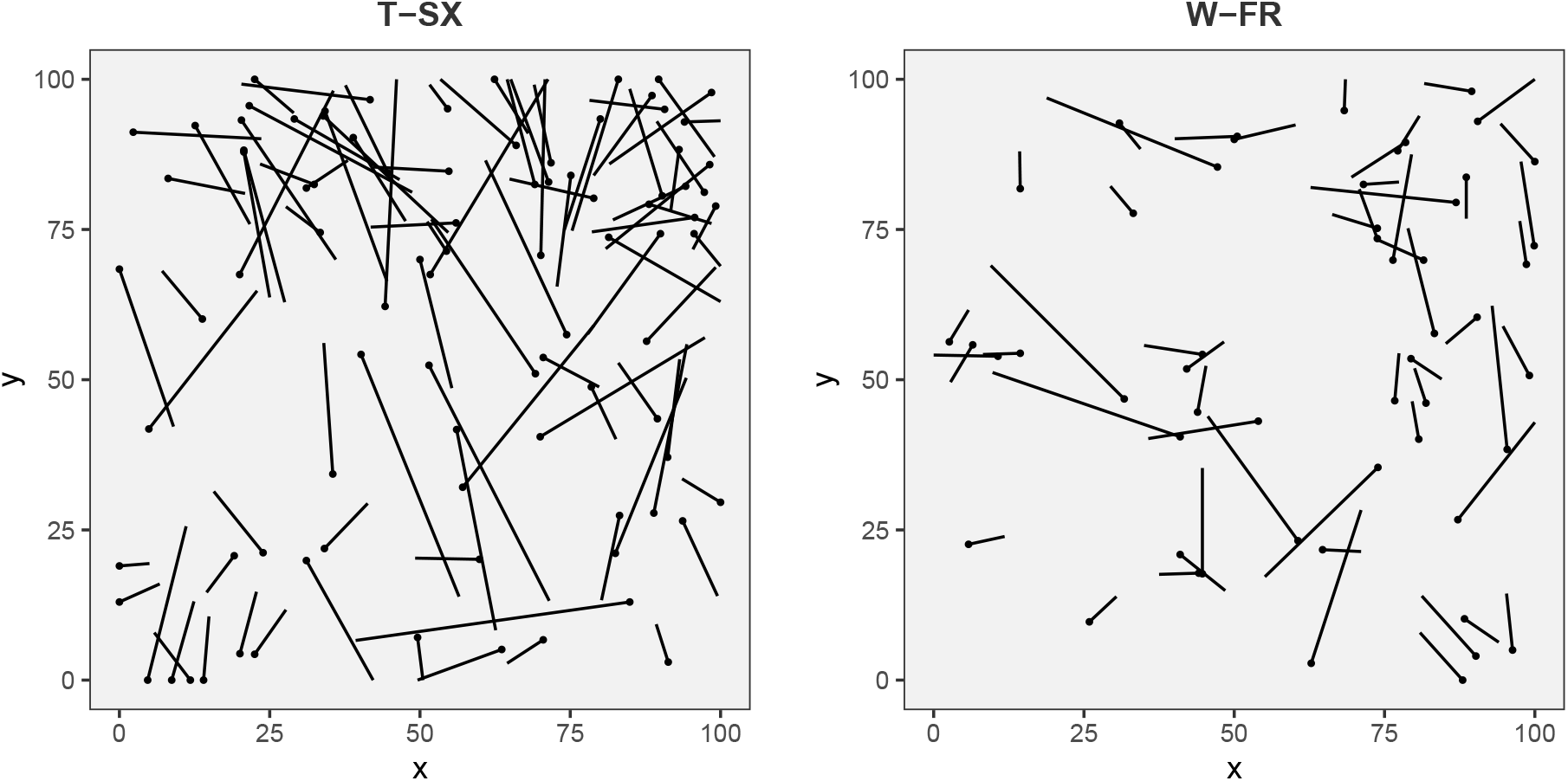
Spatial distribution of logs represented as oriented line-segment patterns. Circles indicate fallen-log bases. Plot dimensions are 100 *m* × 100 *m* for both sites. Site names as per Figure 2.

**Figure 4:**
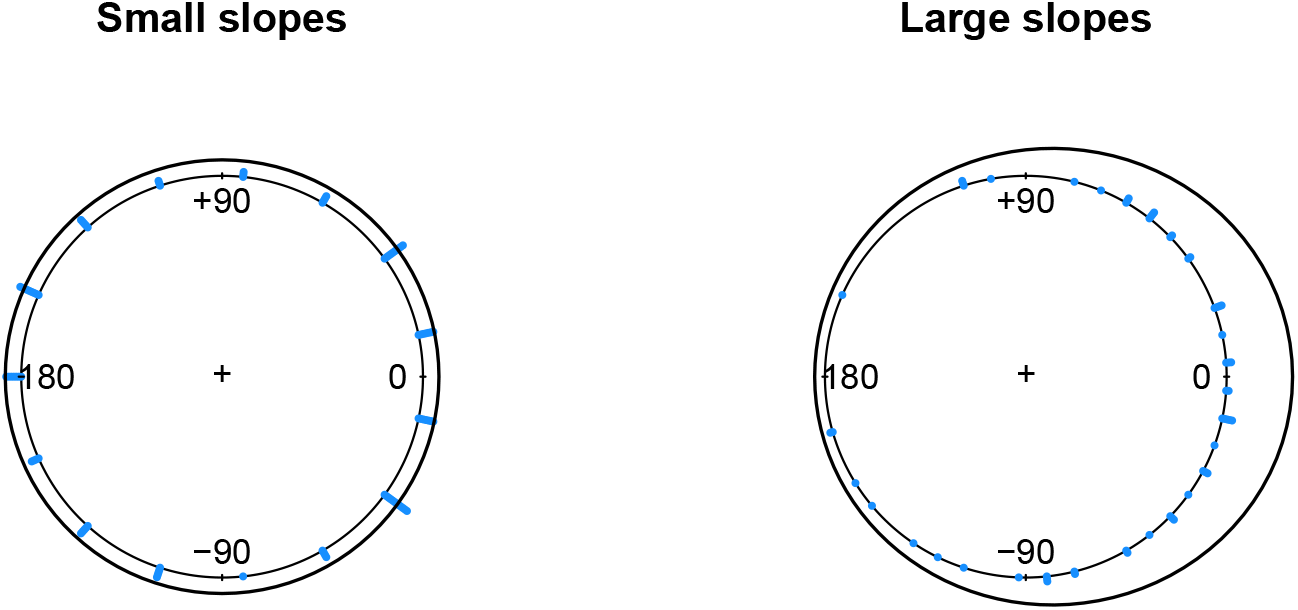
Density plots for local gradient-fitted angular distribution.

**Figure 5:**
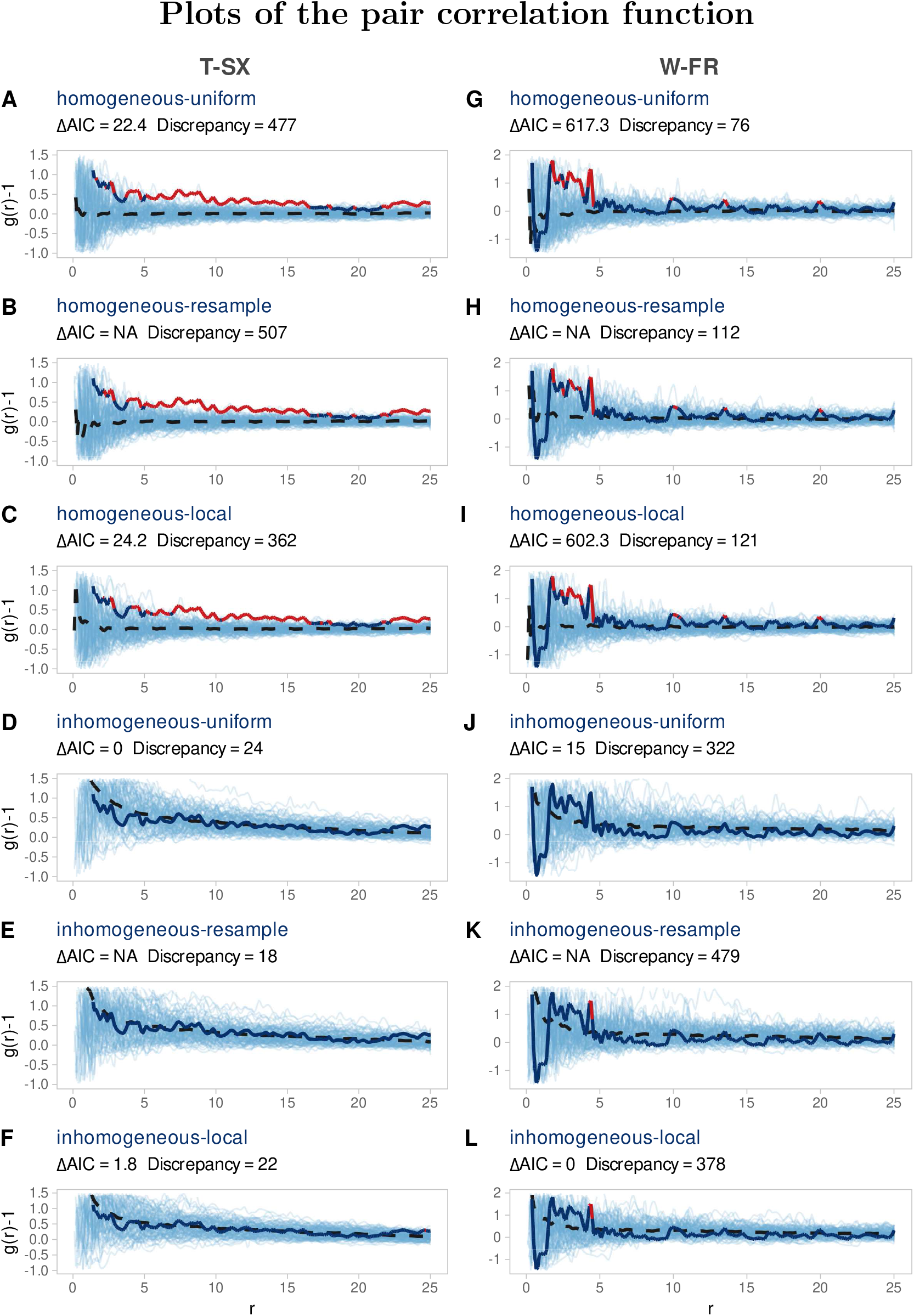
Centred summary functions *g*(*r*) − 1 for both sites. The light blue lines depict the 99 Monte-Carlo simulations, the solid line is the empirical curve, and the dashed line is the mean value of all *g*(*r*) − 1. Viewed as a null-hypothesis test for each model, the progress of the Monte-Carlo *p*-value associated with the rank of the *g*(*r*)-DCLF test statistic (9), computed using *g*(*r*) instead of *L*(*r*), is indicated by the colour of the empirical curve: red if *p* ≤ 0.05, otherwise dark blue.

The discrepancy values are a function of the maximum neighborhood distance *R*, here chosen somewhat arbitrarily, however specific ecological questions may suggest a natural distance scale for which the discrepancy measure becomes more meaningful. There is also the possibility of using the discrepancy, for a chosen value of *R*, to estimate model parameters via minimum-contrast estimation.

All analysis were performed in Program R (R Core Team, 2020). We made extensive use of the package spatstat (Baddeley et al., 2015) which provides basic support for storing line-segment data, and includes an efficient implementation of summary-function and model-fitting algorithms for ordinary point patterns. Estimation of the summary functions for line-segment data is much more computational intensive than that of point patterns. The analysis for this paper, on two forest plots, took 20 hours of computing time, running 20 cores in parallel.

## Results

For both sites, the inhomogeneous models perform substantially better than their homogeneous counterparts, where the apparent ‘clustering’ of logs is best explained by the fitted log-quadratic intensity (Figure 2). This does not allow us to infer that logs are not clustering at all (through, for example, the mechanism of large trees knocking down neighbouring trees as they fall), but it does suggest that any clustering effects scale smoothly with density, since a low-order polynomial surface is sufficient to generate similar summary statistics under model simulation.

The effect of uniform versus non-uniform angular distributions varies between the two sites. For T-SX, with a higher density of fallen logs, the discrepancy is slightly reduced for the non-uniform models, however the AIC score suggests that the additional complexity is unwarranted (Figure 2D-F). In the absence of an AIC score, or a method to calibrate the significance of the *difference* in discrepancy values, it is difficult to assess the relative merits of the re-sampled model. For W-FR, with a sparser distribution of fallen logs, the discrepancy and AIC scores give contradictory results. AIC shows a strong preference for the most complex, inhomogeneous-local model, whereas the discrepancy suggests a better fit is obtained using the inhomogeneous-uniform model. In this instance, given the use of maximum likelihood estimation, AIC is arguably a more ‘natural’ choice of model score, since it estimates predictive log-likelihood (but see ‘Comparison of methods’).

For T-SX, the null hypothesis of homogeneous intensity is rejected (*p* ≤ 0.05) at most distances *r*, independently of the type of angular distribution (Figure 2A-C). For W-FR, the null hypothesis of homogeneous-uniform is rejected at distances close to 5m, however, homogeneous models with non-uniform angular distributions are not rejected.

Although we have demonstrated the use of both model selection and null-hypothesis testing in these two examples, we do not advocate their simultaneous use — in principle, model selection estimates the relative merits of multiple-working hypotheses, placing each on a equal footing (Burnham and Anderson, 2002). However, the use of null-hypothesis testing is well-established in the analysis of spatial point patterns, where a typical analysis involves the selection of a null model in response to both initial inspection of the estimated summary statistic(s) for the data and the specific biological question asked (Wiegand and A. Moloney, 2004). For either method, the use of Monte-Carlo methods to generate plots of simulated summary statistics are a useful form of model validation, where the ability of a selected ‘best’ model to generate plausible data sets can be visually assessed (Gabry et al., 2019).

## Discussion

We have introduced a concrete and accessible set of methods to analyse the spatial patterning of line-segment data. These are based on a natural extension of the familiar (point-pattern) summary function, Ripley’s *K*, and they include simple closed-form expressions for the corresponding estimators and edge correction terms. Using stochastic line-segment processes to model candidate hypotheses, we applied the new methods to a case study of two Australian eucalyptus-forest plots. Indeed, as supported by Buettel et al. (2017) and Kennedy and Spies (2007), we found that treefall is best explained by plot-level spatial heterogeneity. That is presumably unmeasured local-scale factors like (creeklines and gulleys, rock piles, understory trees *<* 10*cm* DBH etc.), which exert the most influence on the final resting place of fallen trees with a plot (Oberle et al., 2015; Buettel et al., 2018; Maser et al., 1998). This suggests that future work might seek to measure these finer-scale, and largely abiotic plot features in combination with the measurements of the living and dead (fallen) woody biomass.

The novel methods and framework we have developed are related to, but also distinct from, various stereological results which are typically presented in integral form and based on the neighborhood of a random point rather than a random segment (Chiu et al., 2013). Although many alternative techniques exist to analyse line-segment data based on the reduction to an associated point pattern (Penttinen and Stoyan, 1989; Cowan, 1979; Eckel et al., 2009), they are less suitable to the application of problems like treefall, as the corresponding measures of inter-point distances do not capture the variation of inter-segment spacing along the length of each segment. Our new summary functions retain this spatial information, thus characterising treefall density within the neighborhood of an entire (random) fallen log. Certain aspects of our methods were nascent in Fortin and Dale (2005, p.62) who proposed a simplified form of the *K*-function for line segments and highlighted the need for further research.

In the development of the summary functions *K*(*r*) and *L*(*r*), we have been careful with the standardisation and transformation of the corresponding estimators. This allows comparability between data sets collected from different sites, which may differ in plot size and sample characteristics, and importantly, between each replicate within sets of simulated data. As a result, we were able to demonstrate the use of Monte-Carlo methods for the comparison of models specified by stochastic line-segment processes.

The definition of a Poisson line-segment process, when characterised by parametric density functions, permits model fitting using maximum (numerical) likelihood estimation, in turn, enabling model comparison within the framework of information-theoretic model selection. In the simplest case, where all models are likelihood-based, a statistical summary function such as *K*(*r*) in not needed for either fitting or selection, although it provides a different perspective on the analysis. For more complex processes, which include interactions such as cluster or inhibition processes, the summary function serves as an important diagnostic tool, enabling the application of hypothesis testing, as well as providing a means to estimate model parameters, via minimum-contrast estimation, when likelihood estimation is intractable (Baddeley et al., 2015, p483).

### Future Directions

In the context of treefall data, given the inert nature of dead logs, interaction models would seem physically implausible, however, bivariate or marked extensions of the *K*-function could help answer ecological questions. This is particularly useful when the inherent attributes of the fallen logs, such as their decomposition status, size (DBH, biomass), or species identity (where possible), are also collected. For example, bivariate analysis of the spatial pattern of small logs (typically saplings and shed branches), in relation to large logs (mostly the trunks of dead trees) could provide insight into the mechanism of treefall. At the technical level, the extension of the methods to include bivariate and marked line-segment patterns should be relatively straightforward. Bivariate forms may even be simpler than univariate, as there is no need to remove the reference line when estimating densities. It is not evident, however, how to extend the definition of the summary function to implicitly account for spatial inhomogeneity; a generalisation of the second-order intensity re-weighted approach due to (Baddeley et al., 2000) would need to solve the problem of attributing a weighting to each segment or possibly to discrete sub-segments. There is currently only one example of a gradient-based inhomogeneous application in the stereological literature (Hahn et al., 1999). An understanding of the interactions between the linear features, as enduring forest ‘logbooks’ (Buettel et al., 2017), can allow us to unpack the impact of legacy disturbance events (wind-throw, fire) and processes (competition) on mortality, diversity, and turn-over in forests (Lassauce et al., 2011).

It would also be of interest to generalise to line-segment patterns, the definition of other traditional summary statistics such as the nearest-neighbour and the spherical-contact distributions; when used in conjunction with second-order statistics, are known to be effective in differentiating between certain types of stochastic processes (Wiegand et al., 2013). Line-segment patterns should also lend themselves to recently developed techniques, such as persistent homology, that exploit the underlying (mathematical) topology of the data, see for example (Robins and Turner, 2016; Pereira and De Mello, 2015; Mecke and Stoyan, 2005). These methods demonstrate resilience to noisy data and increased capacity to identify complex generating processes.

Despite the bias-correction terms in the *K*-function, small edge effects remain present in this study. First, given the relatively large lengths of the logs with respect to the dimensions of the observation window, the estimate 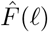 will be biased downward as longer logs are more likely to intercept the boundary and thus be omitted or else truncated. Second, the simulations will underestimate the intensity near the boundary due to the omission of (simulated) logs whose bases lie outside the window yet may have fallen within it. Possible solutions to these problems include simulating over a larger area and using so-called plus-sampling during data collection (Illian, 2008, p.183). The closeness of the mean, centred *L*-function to the horizontal line *y* = 0 for the homogeneous-uniform simulations suggests that these edge effects are minimal, at least in these example forest plots (Fig. 2A,G).

### To Conclude

The present work is an important antecedent to the development of a summary function that incorporates the spatial distributions of both line segments and points simultaneously. In applications for forest ecology, incorporating the statistics of dots and lines into a single analytic framework would allow the relationship between standing, living trees (points) and their dead counterparts (lines), lying on the forest floor, to be examined in new ways. This should provide insights into the role of treefall and coarse-woody debris on the structural patterning of living trees. The hypothesis here would be that the logs have a strong influence on the pattern and structure of the forest both during, and after, the actual treefall event. An understanding of the spatial patterning (and the underlying process) of treefall is critical for the development of integrative models that capture such ecological dynamics and quantitatively describe its role in the generation/maintenance of forest structure and biodiversity.

## Data availability

The data used in this manuscript, together with code to produce the main results will be made available at a figshare digital repository.

## Acknowledgements

This work was funded by the Australian Research Council grant FL160100101.

## Author contributions

All authors conceived the ideas; LY developed the mathematical results, and analysed the data; LY and JB led the writing of the manuscript. All authors contributed critically to the drafts and gave final approval for publication.

## Footnotes

1 Ripley (1976), who showed in general terms that the theoretical properties of a second-order stationary point process may be characterised by the density, λ, and the distribution of inter-point distances.

## Supplementary Materials

### Local gradient-fitted angular distribution

We aggregate the data for both sites and define *θ_r,i_* to be angular difference between the orientation of the *i*th log, *ϕ_log, i_*, and the direction of the (negative) gradient vector at the location of the logs’s base, *ϕ_grad, i_*, that is

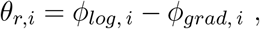

where all angles are given counter-clockwise relative to the *x*-axis.

To identify an appropriate statistical model for *θ*, we follow Pewsey et al. (Circular Statistic in *R*, §6.4), taking as a global model the *Jones-Pewsey* distribution characterised by three parameters: *μ* (location), *κ* (concentration), and *ψ* (shape). This family contains the following set of two-parameter sub-models: wrapped Cauchy (*ψ* = −1), cardiod (*ψ* = 1) and von Mises (*ψ* = 0) distributions.

To perform model selection, we generate a list of candidates comprising the Jones-Pewsey family of distributions, together with the two-parameter wrapped normal distribution and the one-parameter, centred (*μ* = 0) von Mises distribution. Finally, to account for the known dependence of treefall direction on the *magnitude* of the gradient (trees tend to fall downhill), we use a profile log-likelihood approach, to determine a threshold for the magnitude (slope), below which the angular distribution is modelled as uniform rather than gradient dependent. Using AIC, the selected best model is the centred von Mises distribution, characterised by the concentration *κ* = 0.9339, with a slope threshold of 0.21. The angular data and the modelled probability-density functions are shown in Fig 4.

## Elevation and slope plots

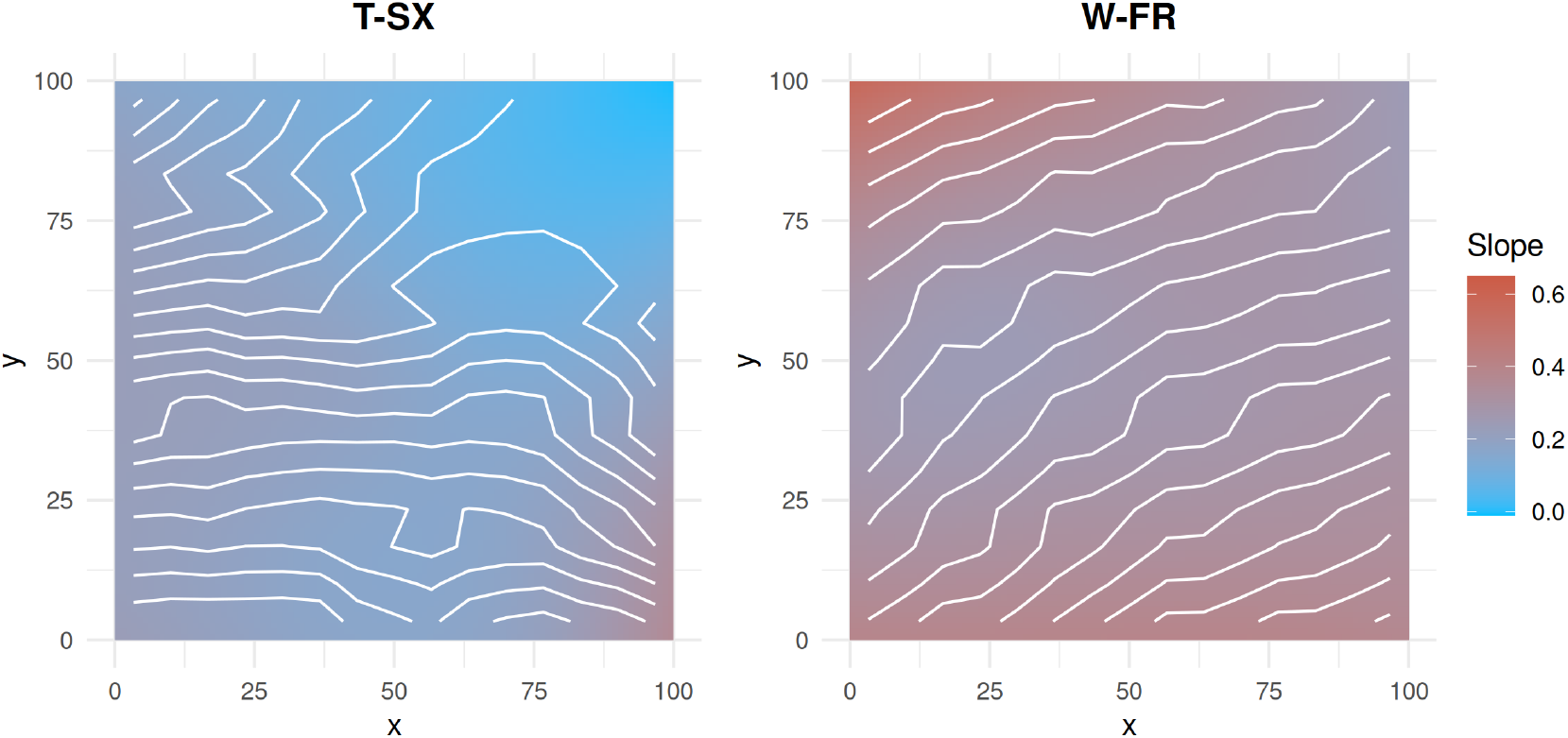

